# Remodeling mechanisms determine size distributions in developing retinal vasculature

**DOI:** 10.1101/2020.06.16.154344

**Authors:** Osamu Iizuka, Shotaro Kawamura, Atsushi Tero, Akiyoshi Uemura, Takashi Miura

**Author notes:** These authors contributed equally to this work (Osamu Iizuka, Shotaro Kawamura).

## Abstract

The development of retinal blood vessels has extensively been used as a model to study vascular pattern formation. To date, various quantitative measurements, such as size distribution have been performed, but the relationship between pattern formation mechanisms and these measurements remains unclear. In the present study, we first quantitatively measured the island size distribution in the retinal vascular network and found that it tended to exhibit an exponential distribution. We were able to recapitulate this distribution pattern in a theoretical model by implementing the stochastic disappearance of vessel segments around arteries could reproduce the observed exponential distribution of islands. Second, we observed that the diameter distribution of the retinal artery segment obeyed a power law;. We theoretically showed that an equal bifurcation branch pattern and Murray’s law could reproduce this pattern. This study demonstrates the utility of examining size distribution for understanding the mechanisms of vascular pattern formation.

## Introduction

The development of the retinal vasculature has extensively been investigated as a model system for vascular pattern formation [1]. In mice, the retinal vasculature is established during the perinatal period. Initially, astroglia migrates from the optic disc region, developing an astroglial meshwork. Next, the endothelial cells migrate from the optic disc region on this preexisting astroglial meshwork, forming the first capillary network beginning at postnatal day 1 (P1) (Fig. 1a). Subsequently, the hyaloid arteries connect to multiple sites within the capillary network near the optic disc, thereby dramatically increasing the blood flow to the network and driving vascular remodeling. Initially, blood with high oxygen concentration runs through the arteries, leading to capillary regression around the arteries. This region of capillary retardation is called the *avascular zone* [2]. Simultaneously, the blood flow causes changes in the arterial segment diameter, resulting in the formation of an arterial vascular tree [3] (Fig. 1b).

**Fig 1.**
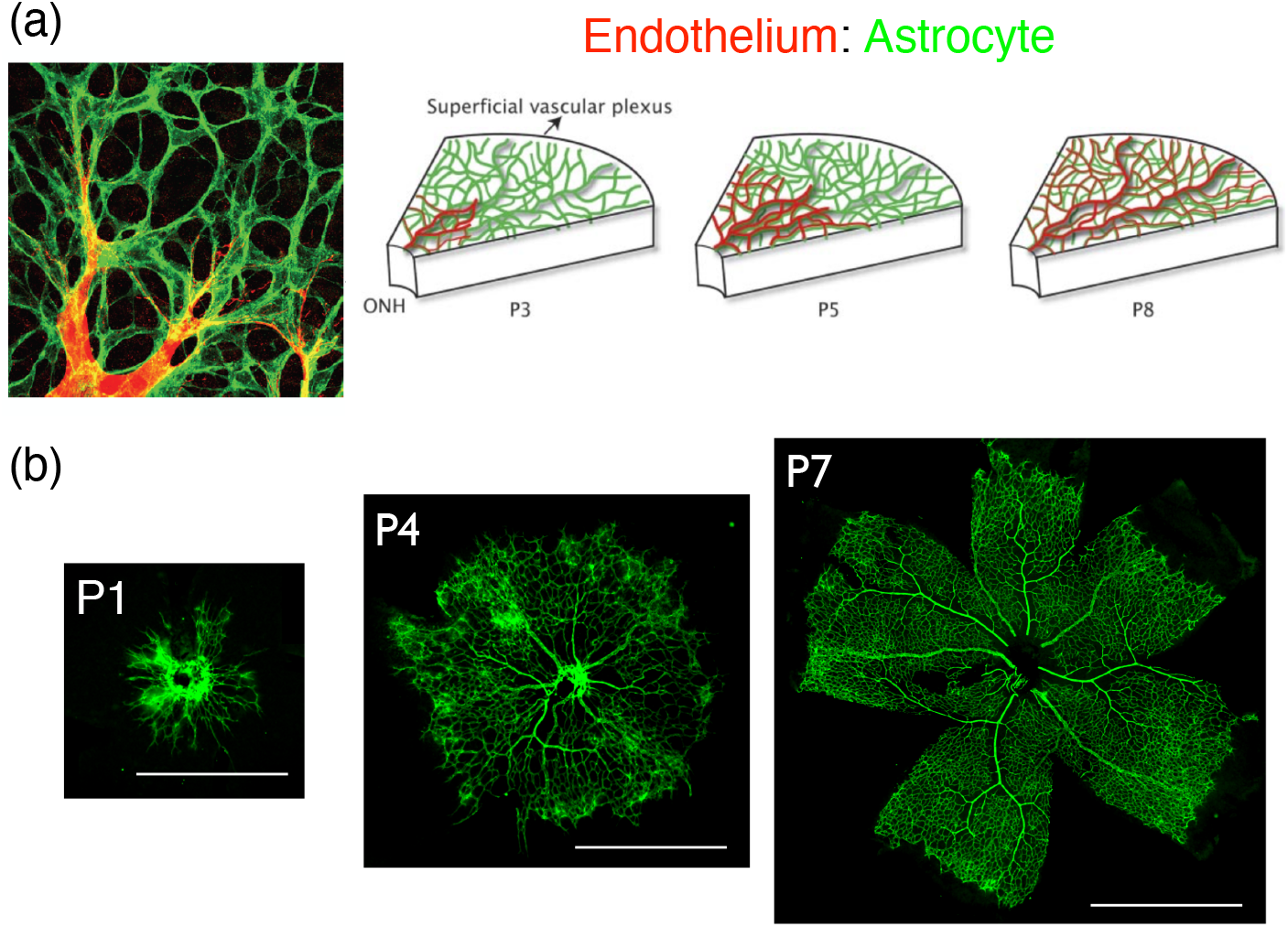
Development of the retinal vasculature in mice. (a) Initially, the astroglial meshwork (green) is generated at the retinal surface. Thereafter, endothelial cells (red) from the optic disc migrate toward the astroglial meshwork, forming the vascular network [1]. (b) With the establishment of blood flow, the arterial diameter gradually changes, forming a branched arterial tree. Scale bars: 1 mm.

Various theoretical models of vascular remodeling have been proposed. Classically, the general relationship between the vascular radius before and after branching has been explained in terms of an energy minimization problem known as Murray’s law [4]. Similarly, the general argument of scaling in various networks has been proposed, which has become an influential concept in physics [5]. More recent studies have reproduced the retinal vasculature remodeling process observed in the avian yolk sac [6] and retinal vasculature [7] using a simple vascular diameter growth rule. Additionally, a large scale computational model that combined the molecular regulation of cell migration, vascular flow, and gas exchange succeeded in reproducing the pattern formation of retinal vasculature [8].

Quantitative measurements of structure can be a useful tool for understanding mechanisms of pattern formation. However, the relationship between pattern formation models and measured quantities has not yet been thoroughly addressed. Previous efforts concentrated on quantifying various aspects of the vascular pattern [9, 10]. For example, the fractal dimension was frequently used as a measure of the vascular pattern [11, 12]. In these studies, the fractal dimension of retinal vasculature was quantitatively measured using the mass-radius or box-count method. However, it is unclear how these quantities are correlated to the pattern formation mechanism. A previous study by [11] asserted that the pattern formation mechanism is based on either diffusion-limited aggregation or Laplacian growth owing to the similarity of the fractal dimension *D* = 1.7. However, based on our current knowledge of the above described retinal vasculature development, this argument is too naive.

In the present study, we combined the above mentioned recent experimental findings, mathematical modeling, and image processing techniques to elucidate the relationship between vasculature size distribution laws and pattern formation mechanisms. First, we focused on the formation of the avascular zone, and experimentally observed that the island size distribution obeyed an exponential law. We formulated a minimal model for the stochastic disappearance of capillaries surrounding the arteries, and theoretically demonstrated that the island size distribution generated by the model exibited the same exponential distribution as was observed experimentally. Second, we focused on arterial remodeling, and experimentally found a power distribution of the radii of the arterial vascular tree. We also confirmed that Murray’s law was established during the development of retinal vasculature. Thereafter we theoretically showed that the combination of bifurcated geometry and Murray’s law could result in power distribution between the segment radius and the number of arterial vascular segments. These results show the utility of examining size distribution for understanding the mechanisms of pattern formation in retinal vasculature.

## Materials and methods

### Immunohistochemistry of mouse retina

All the experiments were undertaken under the permission of the Kyushu University animal experiment committee (A29-036-1). Postnatal day eight (P8) newborn mouse retinas were dissected and incubated in 4 % paraformaldehyde (PFA) overnight. Subsequently, the retinas were washed with PBS and immunostained with 1/1000 anti-alpha-smooth actin antibody (Sigma C6198), 1/100 anti-GFAP antibody (Sigma C9205), or 1/100 anti-CD31 antibody (Abcam ab7388). Endothelial cells were stained with Alexa488-conjugated isolectin GS-IB4 (Labeling & detection, I21411). All images were captured using a Nikon A1 confocal microscope.

### Measurement of the island size distribution

We captured the images of P1 to P8 mouse retinas. For image processing, we used the Fiji software [13]. First, we sharpened the image, transposed it to an 8-bit grayscale image, and manually determined the threshold. Subsequently, we measured the island size using the “particle analysis” command. The distribution was visualized and analyzed using *Mathematica* (Wolfram Research Inc.).

### Measurement of vessel diameter using distance map

We measured vessel diameters by multiplying the distance map and the skeletonized image. First, we made a binarized image of the arterial region using background subtraction and thresholding. After that, we skeletonized the binarized image. Subsequently, we detected the three-way junction points of this skeletonized image using 3 × 3 averaging kernel. Removal of these points from the skeletonized images allowed us to generate a skeletonized image of vessel segments. Following this, we created a distance map using the same binarized image. Finally, we multiplied the skeletonized image of vessel segments and the distance map and measured vessel segment diameters using the “Particle analysis” command. Short segments (< 3 pixels) and extremely thick segments (> 7 pixels) were ignored. Image processing was performed using Fiji software [13], whereas quantification was performed using *Mathematica* Additionally, we assessed the effect of lattice on segment rotation, and the error was < 5 % (S3 Fig).

## Results

### Island size distribution in the vascular network

#### Quantification of the island size distribution in the retinal vasculature

First, we measured the island size distribution in the retinal vasculature (Fig. 2). Vascular structures were detected using the IB4 lectin stain, and the positively stained region was binarized by thresholding. Subsequently, the arterial regions were skeletonized, and the island size distribution was automatically detected using the Fiji software (Fig. 2b, c). The log–linear plot of the histogram showed linearity in size distribution (Fig. 2d). All data points fit well to the linear model, with a exceptions for the smallest island size group in each plot (Fig. 2d, red circles). Moreover, we measured the time course of island size distribution and found that observed linearity was established as the development of the avascular zone (Fig. 2d).

**Fig 2.**
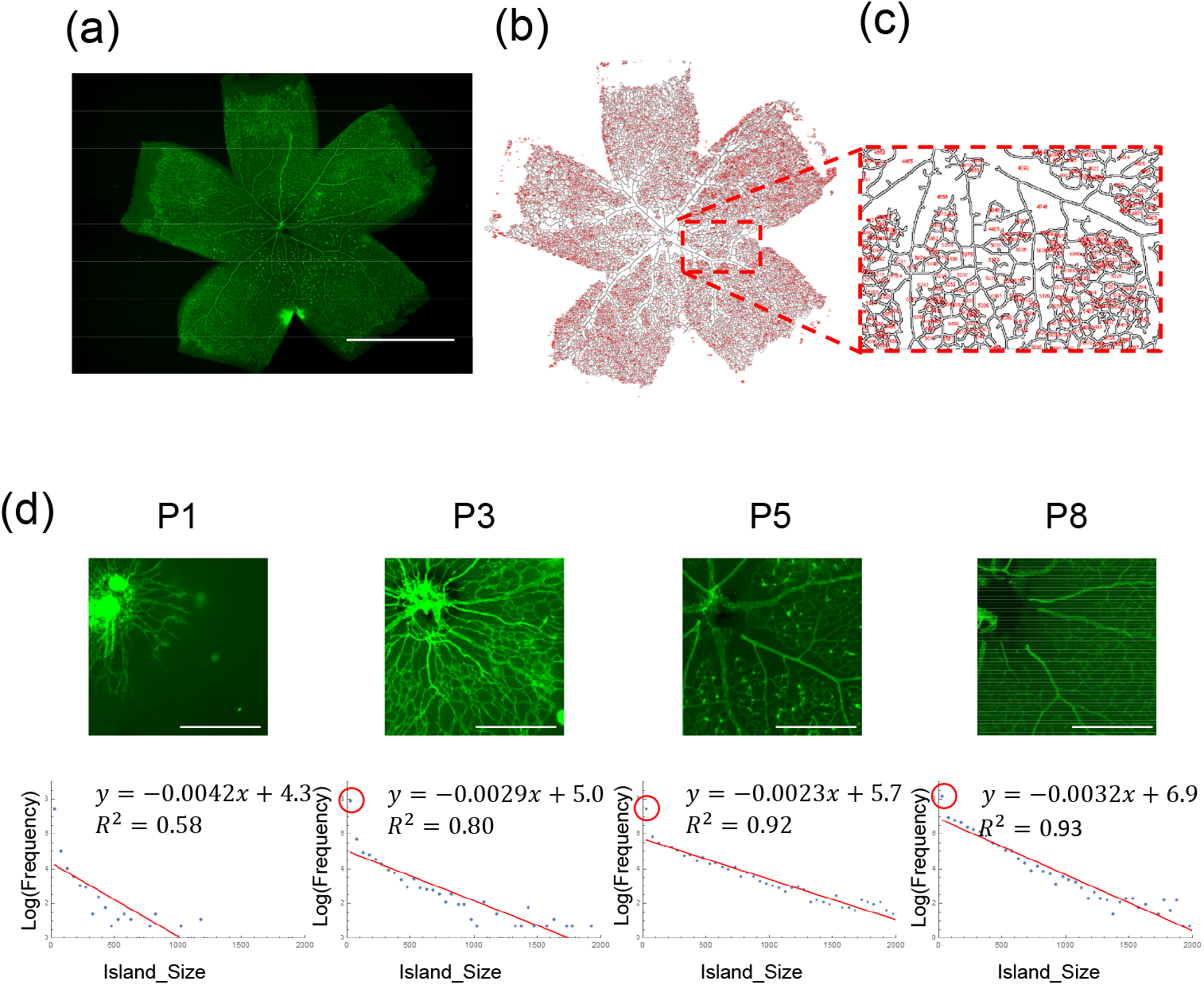
Island size distribution of the retinal endothelial network. (a) P8 mouse retina. Endothelial cells were immunostained with CD31. (b) Automatic measurement of island size distribution. Island size was measured using *Particle Analysis* command in the Fiji software. (c) High magnification view of (b). (d) Time course of island size distribution. At P1, the distribution was scattered. With the progress of development, the distribution gradually became linear in log–*linear* plot, except the smallest size group (red circle). Scale bars: (a) 1 mm, (d) 500 *μ*m.

#### A model to generate large islands in the avascular zone surrounding the arteries

In this section, we propose a simple model that produces islands that exibit exponential size distribution.It has been known that with the establishment of arterial flow, the capillaries near the arteries disappear, owing to the increased oxygen concentration (Fig 3a, [2]). As a result, llarge islands are distributed near arteries close to the optic disc.

**Fig 3.**
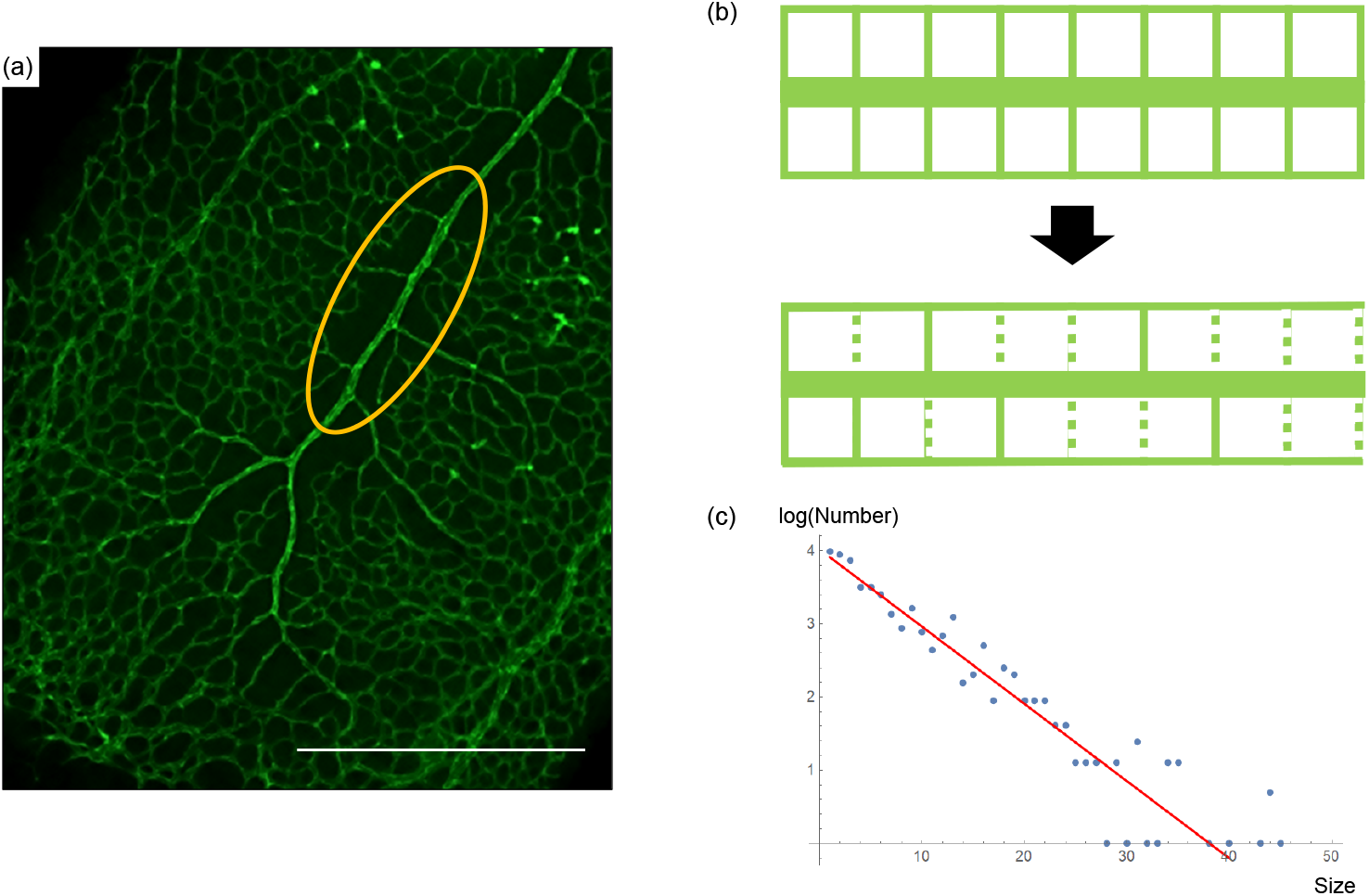
A model of large islands formation in the endothelial network. (a) Large islands in the endothelial network are mainly avascular zones around the arterial tree. This region is generated by the retardation of capillaries in the high oxygen concentration region around the arteries [2]. (b) A minimal model of island formation. At first, the vascular meshwork around an artery is a regular rectangular configuration. As blood flow is established, some of the capillaries adjacent to the arteries disappear by a probability *p* resulting in larger islands of specific size distribution. (c) Numerical simulation of the above model (blue dots) generated size distribution that showed linearity in log–*linear* plot, consistent with the analytical prediction (red line. gradient = log *p*). Scale bar = 50 *μm*.

The region region called the avascular zone. To model this phenomenon, we assumed that the initial vascular meshwork was a regular rectangular lattice, and implemented the well-known fact that the capillaries directly connecting to the arteries disappear in a stochastic manner. In the model, the island distribution is established by the following two-step procedure:

1. A regular mesh of a capillary network is generated.
2. The inter-island capillaries adjacent to the arteries disappear by the probability *p* due to high oxygen concentration.

For simplicity, we assume the initial capillary network is a rectangular lattice. Using this model, we analytically obtained the frequency of islands of size *n* after the stochastic disappearance of capillaries (Table 1). The lattices which are not adjacent to the arteries (*N*_far_) do not change, resulting in the large number of small islands as observed in Fig. 2d (red circles).

**Table 1.** Expected values of number of clusters. *N*_near_, *N*_far_ and *p* represents the initial number of lattices around arteries, the initial number of lattices that are not adjacent to the arteries, and extinction probability of vasculature between islands near arteries respectively.

| Cluster size | Number |
| --- | --- |
| 1 | $N_{\text{near}}(1-p)^2 + N_{\text{far}}$ |
| 2 | $(N_{\text{near}} - 1)(1-p)^2 p$ |
| 3 | $(N_{\text{near}} - 2)(1-p)^2 p^2$ |
| ⋮ | ⋮ |
| $n$ | $(N_{\text{near}} - n + 1)(1-p)^2 p^{n-1}$ |

In this model, the frequency of islands of size *n* was roughly proportional to *p*^*n*−1^, resulting in a linear pattern in the log–linear plot. We also confirmed this tendency numerically (Fig. 3 c). Therefore, this model explains the observed exponential distribution of the island size in the retinal vasculature.

### Power distribution of vascular diameters in the retinal artery network

#### Diameter distribution measurement in the retinal vasculature

Next, we sought to experimentally assess whether the vasculature exhibited Murray’s law and a power distribution between thickness and frequency of vessel segments in the developing mouse retinal arteries. Segment radii were automatically measured using *Mathematica* (Fig. 4). First, we confirmed that the arterial radius could be accurately measured using an *α*-SMA staining (S4 Fig.). To confirm whether Murray’s law holds in the retinal arterial trees, we plotted 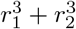 and 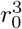, with *r*_0_ indicating the thickest segment. At developmental stages P5 and P6, Murray’s law was not yet established, as indicated by the inequality of the two variables in the plot (Fig. 5b). By P7 and P8, however, the two variables were almost equal, indicating the Murray’s law had been established by the time point. In addition, we plotted the frequency and the thickness of vessel segments in a log–log plot. The power distribution between thickness and frequency was established by stages P7 and P8, as indicated by the linearity of the graph (Fig. 5c).

**Fig 4.**
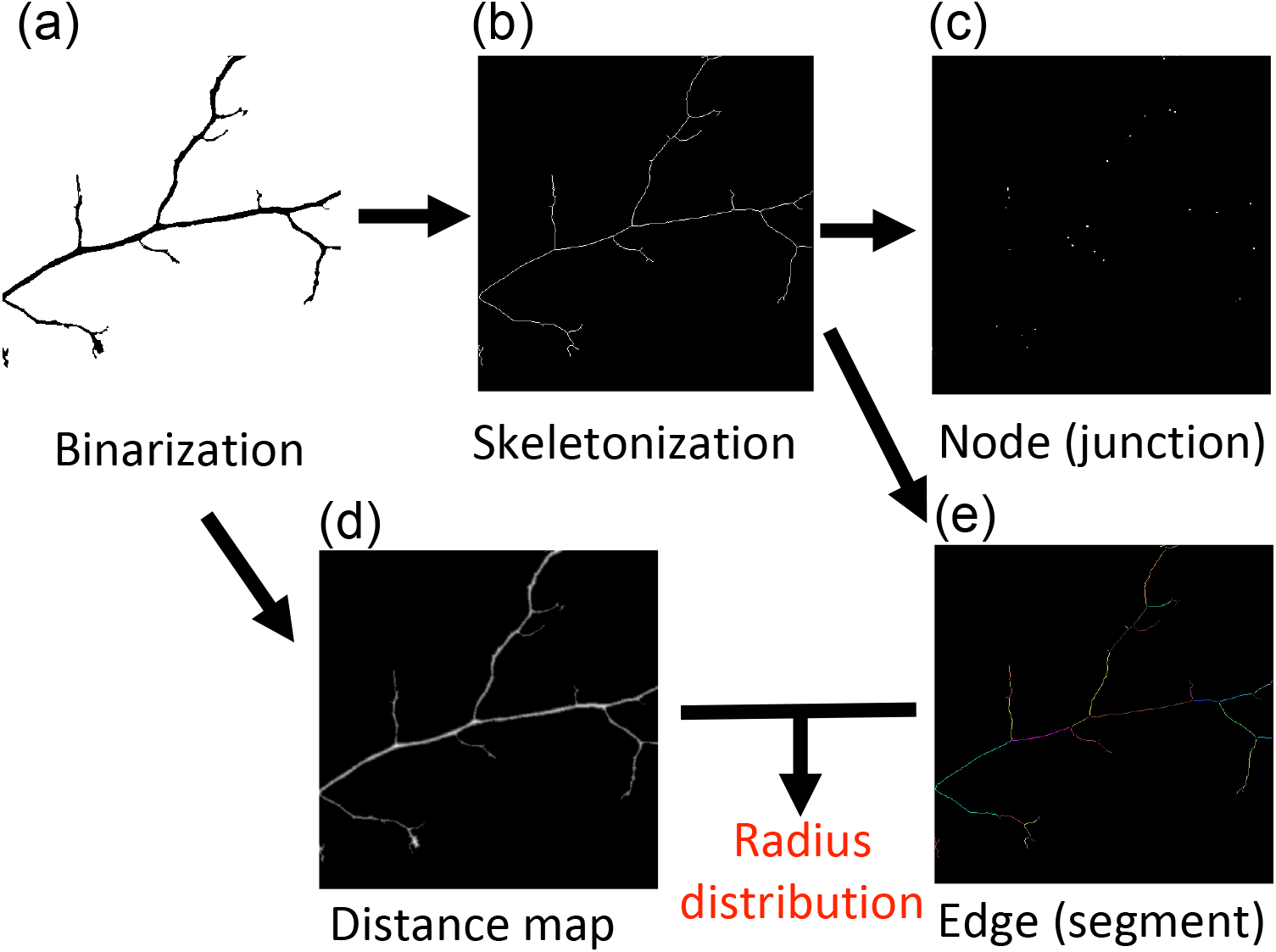
Automatic detection of the segment thickness of the artery region. (a) Images were binarized by a manually determined threshold. (b) The binarized images were skeletonized. (c) Network nodes (junctions) were detected from the skeletonized image. (d) A distance map was created from the binarized image. (e) Network edges (segments) were detected from the binarized image. By multiplying (d) and (e), we obtained radius for each vessel segment.

**Fig 5.**
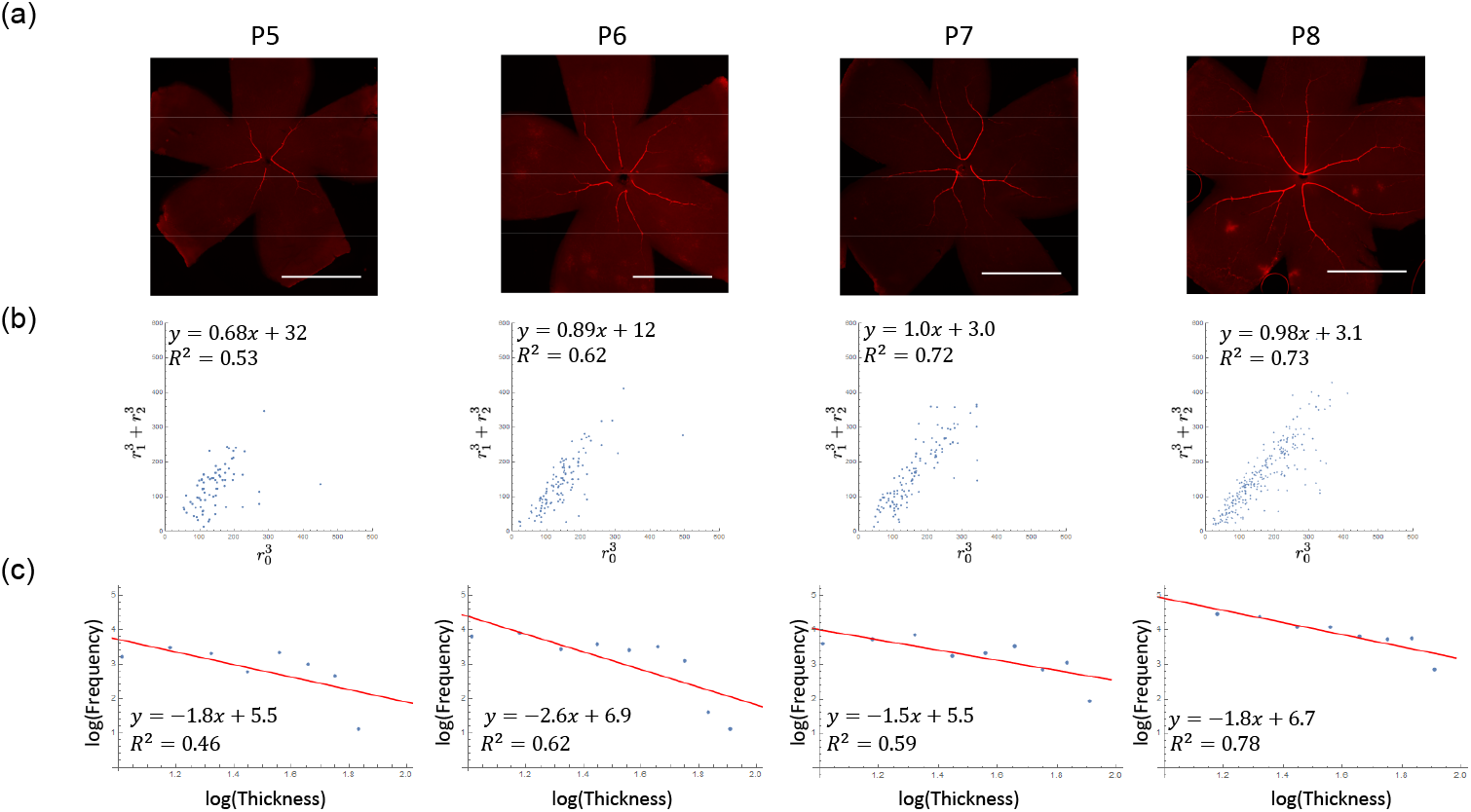
Time course of the relationship between each blood vessel segment thickness and frequency. (a) Retinal images obtained using whole-mount *α*-SMA immunohistochemistry. The tree-like structures of the artery were developed. (b) Relationship between 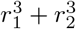 and 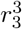. The cubic sums appear to be constant, suggesting that these arteries obey Murray’s law. (c) Log–log plot of frequency versus thickness of each blood vessel segment. Scale bar = 1 mm.

We additionally examined whether the branch angles exhibited Murray’s law, as has previously been predicted [14]. However, the experimentally measured angles did not fit the law (S2 Fig.).

#### Relationship between Murray’s law and vascular branch power distribution

In this section, we show that the combination of Murray’s law and stochastic bifurcation of the retinal artery can result in a power distribution between a vessel segment number and diameter [5, 14, 15]. In Murray’s law ([14], Fig. 6), the following relationship holds at the three-way junctions of blood vessel:

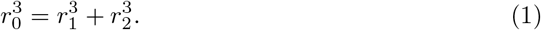

**Fig 6.**
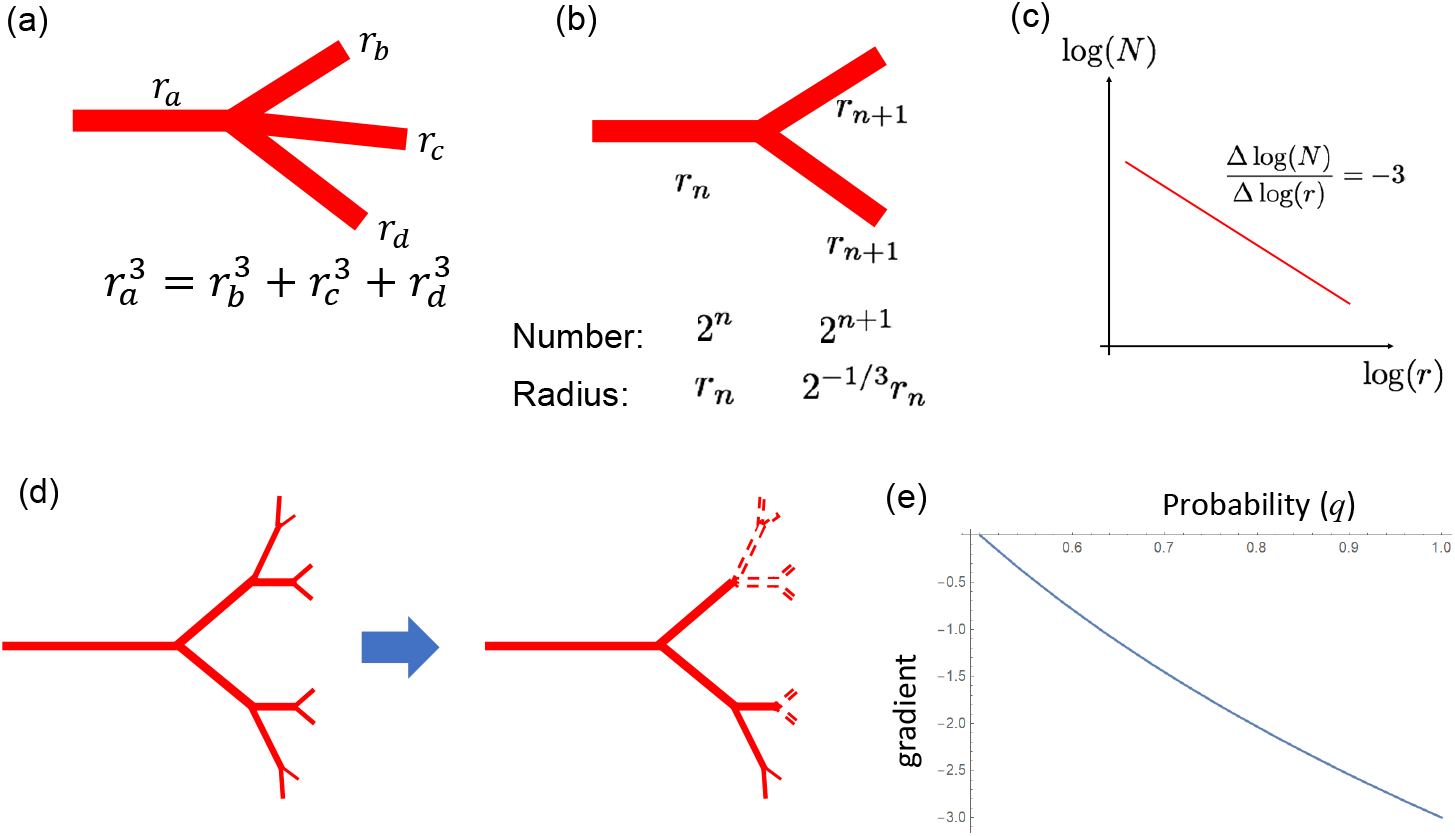
Murray’s law and equal bifurcation leads to scaling of vessel diameter. (a) Murray’s law states that cubic sum of influx *arterial radius* 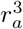 should be equal to the cubic sum of the efflux vessel radii 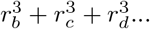 [14]. (b) When the tree structure is generated by equal bifurcation under Murray’s law, it is possible to explicitly obtain the number and radius of the *n* + 1th bifurcation. (c) Each blood vessel segment radius and number should show linearity on a log–log plot, and the gradient should be −3 [16]. (d) Incomplete bifurcation. In the real retina, the arterial tree did not form a complete bifurcation tree. If we assume that the bifurcation occurs at probability *q* the number of thin branches decreases, resulting in a smaller gradient. (e) Relationship between bifurcation probability *q* and gradient. When *q* is less than 1, the gradient is larger than −3.

Assuming that the retinal artery tree consists of repeated equal bifurcations, we obtain two proportional relations one between the mother vessel radius *r_n_* and the sister vessel radius *r*_*n*+1_, and the other between the number of mother vessels *N_n_* and that of sister vessels *N*_*n*+1_. Therefore, the number and radius should obey a power law. With the progression to the next generation, the number of *n* generations doubles from that of *n* − 1th generation, and the radius becomes 2^−1/3^ times that of the former generation (Fig. 6b). Therefore, when drawing a log–log plot, the gradient becomes −3 (Fig. 6c). The establishment of Murray’s law can be explained by setting an appropriate vessel diameter growth function (supporting text 1, S1 Fig.).

With a probabilistic equal bifurcation, the log–log plot exhibits a gentle gradient (Fig. 6d). When *q* is the probability of generating the bifurcation, we can estimate the gradient of the plot accompanying a change of *q* as follows:

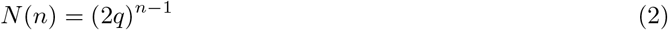

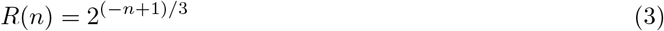

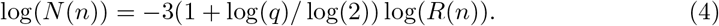

The gradient is expressed as −3(1 + log(*q*)/log(2)). As *q* decreases, the gradient becomes larger than −3 (Fig. 6e), which can explain the gradient observed in Fig. 5.

## Discussion

In the present study, we evaluated the relationship between size distribution laws and pattern formation mechanisms. Among the size distribution laws, power distribution could be correlated to the fractal dimension. Following the original study by [11], several similar investigations were performed focusing on a fractal dimension of retinal vasculature as a diagnostic tool [12, 17–22]. To date, only a small number of studies have correlated size distribution with fractal pattern formation mechanisms, which were not based on the current understanding of retinal vasculature development [23].

*p* and *q* which are experimentally obtained, can be used as a measure for defective pattern formation in the retinal vasculature. For example, *p* can be correlated to the degree of oxygen supply during retina development. *q* can be interpreted as a measure for vascular remodeling by flow and arterial differentiation. It would be intriguing to use these parameters to investigate the shape change induced by various factors such as oxygen-induced retinopathy or an absence of pericytes [24].

A previous report has confirmed the prediction of Murray’s law by manually measuring pig coronary arteries [9]. A modification of Murray’s law has been proposed by [25], which considers the vessel wall metabolic cost as follows:

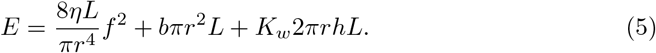

In this equation, *K_w_* represents the metabolic constant, and *h* indicates the vessel wall thickness. In this case, when 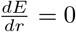 we obtain

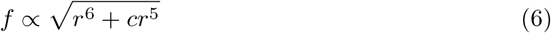

where *c* ∝ *h* However, if this modification holds, we should not observe scaling in vessel segment diameter distribution. We did not need this modification, possibly because vessel wall thickness *h* was very small, and *c* was negligible in our case (S4 Fig. b).

The biological mechanisms of endothelial regression by high oxygen concentration and of vessel diameter change by flow have been well studied. High oxygen concentration results in the inhibition of vascular endothelial growth factor (VEGF) via hypoxia-inducible factor (HIF) [26]. As a result, capillaries are not maintained due to the lack of VEGF. In addition, it has been shown that vessel diameters are influenced by blood flow [27] and pericyte activity [28, 29]. Recent studies have suggested that shear stress regulates endothelial and smooth muscle cell signaling. This signaling is mediated by several factors, including nitric oxide (NO), prostaglandin I-2 (PGI-2), platelet-derived growth factor (PDGF-BB), transforming growth factor *β*1 (TGF-*β*1), and microRNAs (miRs) [30]. There are both fast (NO and PGI-2) and slow (PDGF-BB, TGF-*β*1, and miR126) factors, which are used for communication between the endothelial and smooth muscle cells [31, 32]. Owing to our interest in this structural change, we focused on the latter factors. PDGF-BB and TGF-*β*1 are produced by endothelial cells under low shear stress. Functionally, PDGF-BB activates smooth muscle proliferation, migration, and contraction, whereas TGF-*β*1 activates smooth muscle cell differentiation. Additionally, miR126, which is produced by endothelial cells under laminar shear stress, activates smooth muscle cell proliferation. In future studies, these slow factors can be used to assess the relationship between flow and diameters experimentally. Pericytes are also known to be involved in the retinal vasculature remodeling process. It has been shown that pericytes play a role in modulating the extracellular matrix degradation, production, and assembly via an interaction with endothelial cells [28, 29]. Manipulation of pattern formation and size distribution using these molecules would be an interesting future goal.

## Acknowledgments

The authors want to acknowledge Hiroshi Kori for helpful discussions and insights. This work was financially supported by JSPS Kakenhi (Grant No. 15KT0018) and JST CREST (Grant No. JPMJCR14W4).

